# Using weapons instead of perfume – chemical association strategies of the myrmecophilous bug *Scolopostethus pacificus* (Rhyparochromidae)

**DOI:** 10.1101/2020.12.08.412577

**Authors:** Adrian Brückner

## Abstract

A vast diversity of parasites associate with ants. Living in and around ant nests these organisms must overcome ant colony defenses. As ant defensive behavior is mainly mediated by species-specific cuticular hydrocarbons (CHCs) or alarm pheromones, ant-associated parasites can either crack their hosts chemical communication code by modifying their own CHC-profiles or use pro-active strategies like chemical weaponry for distraction and repellency. While the chemical nature of ant-parasite interactions has been intensively studied for highly host specific parasites, the chemical-deceptive strategies of the rather rare ant-resembling Heteropterans are unknown. To gain insight into this system, I studied the bug *Scolopostethus pacificus* (Barber 1918) which can be found near the nests of the ecologically dominant and aggressive velvety tree ant (*Liometopum occidentale*, Emery 1895). Using behavioral, chemical and molecular approaches I disentangled the relationship of *S. pasificus* and its host ant. Chemical profiling of the bug and the ant revealed that the bug does not make use of CHC insignificance or mimicry, but instead uses a cocktail of volatile compounds released from its metathoracic glands that likely moderates encounters with its aggressive host. Feeding trials with armed and artificially disarmed bugs revealed a defensive function of the gland exudates. Targeted molecular gut barcoding showed that *S. pasificus* does not feed on *L. occidentale*. These results suggest that chemical weaponry, rather than a chemical code-cracking CHC matching or chemical insignificance, enables *S. pasificus* to get along with and live in close proximity to its host ant.

## Introduction

Parasites use a variety of chemical strategies to deceive other species and mask the true nature of their interaction (Bagnères and Lorenzi 2010). Numerous examples of deception and trickery have been described in parasites of social insects (i.e. ants, termites, bees and some wasps) (Dettner and Liepert 1994; Guillem et al. 2014; Parmentier 2020). For those parasites succeeding in overcoming social insect colonies’ defense, the companionship with their host guarantees protection from the environment as well as access to resources (Kistner 1982; Parker 2016).

Ant parasites must overcome a nestmate recognition system based on a species- and colony-specific blend of long-chained cuticular hydrocarbons (CHCs). Like most other insects, ants secrete CHCs onto their epicuticular exoskeleton, primarily as a protection against water loss (Blomquist and Bagnères 2010; Howard and Blomquist 2005; von Beeren et al. 2012). Additionally, ants use these CHCs in a process called label-template matching to recognize nestmates and root out opponents (e.g. predators, other competing ants, or parasites) (Blomquist and Bagnères 2010; Martin and Drijfhout 2009). Upon discovery, these colony-intruding organisms are often attacked and killed. Many social insect parasites outmaneuver the chemical alarm system of their hosts using chemical-deceptive strategies like *de novo* chemical mimicry, acquired chemical mimicry, chemical insignificance and/or chemical weaponry (Akino 2008; von Beeren et al. 2012). Chemical mimicry and insignificance are very well studied tactics of ant colony parasites: many arthropods that successfully integrate themselves into heavily armed ant nests match their host CHC profiles either actively *via* biosynthesis of similar CHCs or passively by acquiring CHCs from their host’s cuticle through occasional contact and/or specific grooming behavior (Akino 2008; Bagnères and Lorenzi 2010; Lenoir et al. 2001; von Beeren et al. 2011). Other parasites have a greatly attenuated CHC layer on their cuticle surface allowing them to be chemically invisible to their host. This can be achieved with a quantitative reduction in all CHCs in the profile or the removal of certain key compounds necessary for nestmate recognition (Dettner and Liepert 1994; Uboni et al. 2012; von Beeren et al. 2012). This strategy of chemical insignificance might often be used in parasites’ pre-colony integration stage to initially infiltrate the host (Uboni et al. 2012). Besides CHC-based trickery, parasites can also take a more aggressive approach and use defensive allomones (chemical weaponry) to counteract the host’s own defensive behaviors or disrupt the nestmate recognition (Akino 2008). This strategy has been predominantly described in interactions between ants and socially parasitic ants or paper wasps parasitizing ants (D’Ettorre et al. 2000; Neupert et al. 2018; Post and Jeanne 1981).

Many highly specialized “myrmecophiles” – here defined as parasites whose life history depends at least in some aspects on their social interaction with the ant colony without returning obvious benefits (Parker 2016) – that are obligately associated to their host, use chemical mimicry and display a closely matching CHC profile (Akino 2008; Bagnères and Lorenzi 2010; von Beeren et al. 2018). Such parasites are often socially integrated into the ant society and are treated as nestmates. This manifests in mouth-to-mouth food exchange with ant workers, in being carried by workers, and/or in achieving access to sensitive parts of the nest for instance the brood chambers (Hölldobler and Wilson 1990; Kistner 1981; Parker 2016). Many obligate myrmecophiles exhibit specific morphological adaptations to the life with ants (Kistner 1981; Kistner 1982). For instance, some staphylinid beetles even bear close morphological resemblance to and share similar coloration as their host (Maruyama and Parker 2017). Most obligate myrmecophiles usually feed on brood or steal food and in some cases live as ectoparasites (Kronauer and Pierce 2011; Wasmann 1894). On the other hand, large numbers of ant parasites live as unspecialized associates that are not accepted as nestmates and provoke colony defense at least at a certain level (Parmentier et al. 2014; Parmentier et al. 2017). Facultative myrmecophiles usually do not share distinct morphological features and seem to use chemical insignificance to sneak into their hosts’ nests to prey on brood, food collected by the colony or even hunt other myrmecophiles (Kistner 1982; Parker 2016; Parmentier et al. 2017). Other facultative parasites do not employ CHC mediated disguises and most likely have to rely on other strategies such as chemical weaponry or behavioral evasion strategies (Akino 2008; Lenoir et al. 2001; Parmentier et al. 2017).

The strategies used by facultative parasites for dealing with ant aggression are currently understudied. The current study addresses the chemical-deceptive strategies of *Scolopostethus pacificus* (Barber 1918) and the nature of its association with the velvety tree ant *Liometopum occidentale* (Emery 1895). Large aggregations of *S. pacificus* – an ant-resembling bug species (Rhyparochromidae) – have been repeatedly found in close proximity of *L. occidentale* nests in Southern California and both species share a strikingly similar color palette (Fig 1A). The velvety tree ant is an ecologically dominant ant species native to the Pacific West of the United States (California, Oregon, South Washington) and Northern Mexico, that hosts a great diversity of myrmecophiles (Danoff-Burg 1994; Danoff-Burg 2002; Hoey-Chamberlain et al. 2013). *Scolopostethus pacificus* does not seem to be a close associate with *L. occidentale* as the bug has been found in leaf litter samples, but has never been found in the actual nest material of the ants (Barber 1918; Larson and Scudder 2018) and appears to be recognized and attacked by the ants (personal observation).

**Figure 1.**
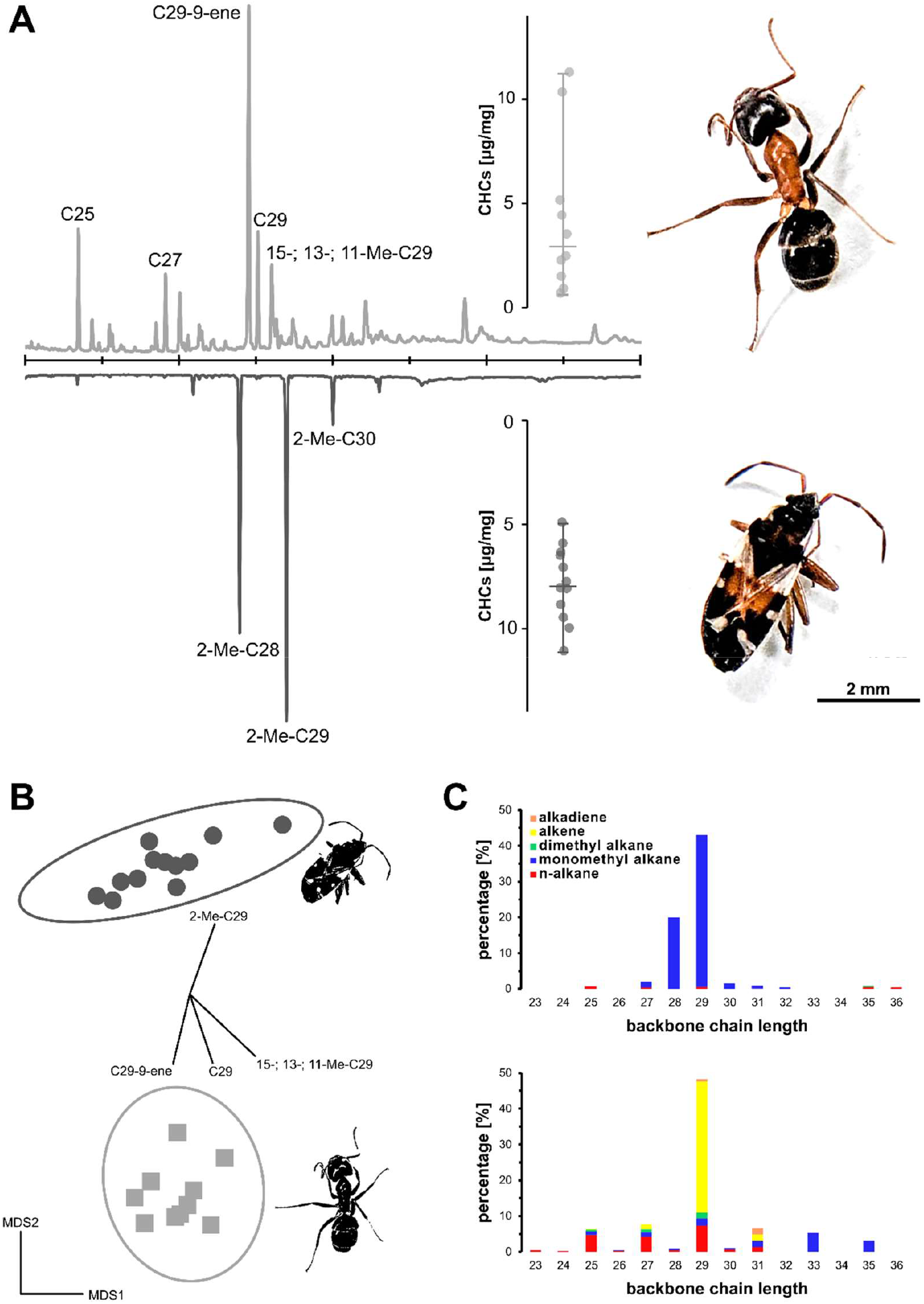
No CHC profile similarities between bug parasite *Scolopostethus pacificus* and the ant host *Liometopum occidentale*. (A) Velvety tree ants and bugs share a similar color palette. Example GC traces for both ants (top) and bugs (bottom) show that there is nearly no overlap in CHC composition. Main CHCs are labeled and a detailed list can be found in Table 1. The amount of CHCs [μg] for both insects (dot plot inserts) were also similar (see results); horizontal bars represent the min, max and median. Ant and bug images are at the same scale. (B) Non-linear multidimensional scaling (NMDS; 2D-stress= 0.12) plot of CHC compositional data based on Bray-Curtis similarities. Each data point represents an individual sample, ordination ellipses represent 95% confidential space around the group’s centroids. CHCs that majorly contributed to data separation are mapped onto the ordination as vectors. Dark-grey dots represent bugs and light-grey squares ants. (C) Differences between CHC profiles of bugs (top) and ants (bottom); plots show the mean distribution of different substance classes per chain length.

Hence, I hypothesized that *S. pacificus* does not use chemical mimicry or insignificance to remain in close contact to *L. occidentale* ants, but instead uses chemical weaponry based on its metathoracic scent gland (MTG) – a chemical defense system known from other Heteropterans (Aldrich 1988). I further expected that similar to other facultative myrmecophiles, *S. pacificus* might either prey on the ants and their brood or benefit from protection against predators – either directly by the ants, or indirectly because of the matching cuticle patterns and coloration (Batesian mimicry). To test my hypotheses, I used a combined approach of chemical analysis of CHCs and MTG compounds, survival assays and targeted molecular gut-content analysis.

## Materials and Methods

### Collection

Individuals of both species, *Scolopostethus pacificus* and *Liometopum occidentale*, were collected from two ant nest sites in close proximity on July 2018 at Rubio Canyon, Altadena, CA (34°12’19.7”N 118°07’01.7”W). Only forager ants that were close to the nest entrance were collected, while bugs similar in size to the ants were picked from aggregations of specimens next to the entrance. Animals were transferred to the laboratory in a cooling bag and directly used for chemical analysis, bioassays or gut-content analysis. Additionally, ant nest material was collected and frozen in the laboratory.

### Chemical analysis

To identify volatile compounds that are released by *S. pacificus* upon disturbance, I used an established SPME sampling method (Brückner et al. 2018): Individuals were squeezed and quickly placed into individual GC vials (glass, 12 x 32 mm, sealed with an 11-mm Teflon-lined septum) and released volatiles were immediately sampled with a 65 µm polydimethylsiloxane/divinylbenzene fiber from Supelco (Sigma-Aldrich; St. Louis, MO). For compound desorption the fiber was placed in the injector port for 1 min operating at 230°C and the GC/MS run was carried out as outlined below. After each trial the fiber was baked for 30 min at 230°C. SPME-GC/MS runs were used to identify the fraction of volatile metathoracic scent gland (MTG) compounds (see e.g. Krajicek et al. 2016), but for quantification of these I used whole body extraction in 50 μl hexane with internal standard for 10 min. For the analysis of cuticular hydrocarbon (CHCs) I submersed freeze-killed ants and bugs in 50 μl hexane with an internal standard (see below) for 10 min.

A GCMS-QP2020 gas chromatography/mass-spectrometry system (Shimadzu, Kyōto, Japan) equipped with a ZB-5MS fused silica capillary column (30 m x 0.25 mm ID, df= 0.25 μm) from Phenomenex (Torrance, CA, USA) was used for chemical profiling. Crude hexane sample aliquots (1 μl) were injected using an AOC-20i autosampler system from Shimadzu, into split/splitless-injector operated in splitless-mode at a temperature of 310°C. Helium was used as the carrier-gas with a constant flow rate of 2.13 ml/min. The chromatographic conditions were as follows: The column temperature at the start was 40°C with a 1-minute hold after which the temperature was initially increased 30°C/min to 250°C and further increased 50°C/min to a final temperature of 320°C and held for 5 minutes. Electron impact ionization spectra were recorded at 70 eV ion source voltage, with a scan rate of 0.2 scans/sec from *m/z* 40 to 650. The ion source of the mass spectrometer and the transfer line were kept at 230°C and 320°C, respectively. Semi-quantification of CHCs and gland compounds was based on the internal C18 (100 ng/ul) or C10 (150 ng/ul) standards, respectively. I integrated the chromatograms manually using LabSolutions Postrun Analysis (Shimadzu, Kyōto, Japan) and aligned the compounds and their respective TICs in Microsoft Excel (version 1908). I quantified the ion abundance and calculated the absolute amounts of compounds in microgram (µg) based on the internal standards, as well as the relative composition of individual compounds compared to the total ion abundance. Both ants and bugs were dried until constant weight and measured on a fine scale. Dry weights were used to standardize the CHC amounts and to calculate the CHC/dry weight (µg/mg).

MTG compounds were identified based on the *m/z* fragmentation patterns and retention times. Additionally, the identities of (2*E*)-hex-2-enal, isoamyl acetate, 3-methyl-but-2-enyl acetate and (2*E*)-hex-2-enyl acetate were confirmed with analytical standards from Sigma-Aldrich (St. Louis, MO). For the analysis of CHCs, all non-hydrocarbon substances as well as CHCs were excluded if their average abundance was consistently below 0.1% of the total CHC abundance. CHCs were identified using diagnostic ions and retention indices calculated based on a standard series of n-alkanes [Carlson et al. (1998); **Table 1**]. To identify double bond positions in alkenes and alkadienes, I performed methylthiolation of the carbon-carbon double bonds using DMDS (dimethyl-disulfide, Sigma-Aldrich, St. Louis, MO) according to Carlson et al. (1989). The configuration of double bonds was not determined.

### Disarming protocol and survival assay

To test the hypothesis that the MTG secretion of adult *S. pacificus* serves a defensive function, I measured the survival of armed vs. unarmed bugs against *L. occidentale* ants. Bugs were disarmed following the protocols of Duffey and Scudder (1974) and Krall et al. (1999), briefly; insects were gently squeezed with a pair of tweezers multiple times along the bilateral axis of the thorax, washed in 0.1x PBST and rinsed in water. This procedure was repeated three times over 1h and bugs were allowed to recover for 1h after the last squeezing. In total, 55 out of 61 bugs survived the disarming protocol and 15 insects were directly submersed in 50 μl hexane with internal standard for 10 min to extract MTG compounds. In addition, 16 untreated, armed bugs were also extracted as control and all gland traces were profiled via GC/MS (see above).

For one survival assay trial, five *L. occidentale* were put into small plastic boxes (105×105×80 mm) with a piece of nest material (**supplement Fig S1**) and slightly wet tissue paper at the bottom, resembling a more realistic nest environment, triggering colony defense (e.g., Kleineidam et al. 2017). Then, a single bug was introduced per box and the assay run over 24h in a temperature-controlled room at ∼21°C. In total, I used 20 armed and 20 disarmed adult *S. pacificus* and recorded their survival by the end of the trial period. Armed bugs that survived the bioassay trial were finally extracted in 50 µl hexane with internal standard for 10 min to profile their MTG secretion. Additionally, 20 disarmed bugs were placed in similar containers without ants, to record a baseline mortality rate.

### Targeted molecular gut-content analysis

For the molecular analysis of the bugs’ gut content, reference specimens of both *S. pacificus* and *L. occidentale* were directly frozen. For gut-content PCR, five guts (4 replicates, 20 guts in total) of *S. pacificus* were dissected in ice-cold 1xPBS and pooled. Gut-free bug bodies were pooled and frozen, too. DNA from the whole specimens, bodies only and the respective guts was extracted with the Zymo Research Quick-DNA Miniprep Plus Kit. DNA extracts of the specimens were used in PCR reactions (initial denaturation 2 min at 95°C; 35 cycles of 0.5 min at 95°C; 1 min at 58°C, 1 min at 72°C; final extension for 5 min at 72°C and final indefinite hold at 4°C) using ITS2 primers [**supplementary Table S1**; Ji et al. (2003)] with GoTaq Green (12.5 µl master mix, 8.5 µl nuclease-free water, 2 ul 10 µM mixture of F/R primer, 2 µl template). A volume (5 µL) of the reaction was then checked for successful amplification on a 1% agarose with the 1 kb DNA Ladder from New England Biolabs and afterwards the remaining PCR product was purified using the Monarch® PCR & DNA Cleanup Kit (New England Biolabs; Ipswich, MA). Purified products for *S. pacificus* and *L. occidentale* were sent for Sanger sequencing with Laragen Inc. (Culver City, CA). Sequence data were aligned and used to design two ant specific primers (Loc1 and Loc2, sequences see **supplementary Table S1**) to evaluate whether the bugs consumed ants. Finally, these new primer sets were used in a PCR reaction using the bug-body DNA, bug-gut DNA and control ant DNA as template following the previously described method (see above). As a positive control guts of ten *Platyusa sonomae* beetles (2 guts per replicate), which had been fed with ants were dissected and subjected to targeted gut-content PCR with Loc1 and Loc2 as described above. Additionally, all remaining parts of the beetle’s bodies were DNA extracted and used for standard ITS2 primer amplification. PCR products were again checked on 1% agarose gels, purified and sent for Sanger sequencing.

### Statistics

All statistics were performed with PAST 4.02 (Hammer et al. 2001) and R 3.6.1 (R_Core_Team 2019).

I assessed the differences in CHC composition between bugs and ants by first calculating a Bray-Curtis similarity (BCS) matrix based on the compositional data of each individual, second visualizing the matrix using non-linear multidimensional scaling (NMDS) plots and third performing a PERMANOVA on the BCS matrix using species as fixed effect (Brückner and Heethoff 2017). To standardize the amounts of CHCs between bugs and ants, I dried the specimens to obtain their dry weight and calculate the CHCs in µg/mg and finally statistically assess significance with a Mann-Whitney-U test.

The MTG composition was visualized using a heat-coded matrix plot. I used Mann-Whitney-U tests to compare MTG secretion of armed vs artificially disarmed bugs and armed vs. bugs that survived the bioassay. Type I error accumulation for this analysis was corrected using the false discovery rate approach (Benjamini and Hochberg 1995). To test the defensive function of the MTG and assess bug survival rates, I used the cbind() command in R to create a binomial response variable (“alive” or “dead” for each sample). This variable was then used in a generalized linear model (GLM) with a binomial distribution and armed/disarmed as fixed effect.

## Results

### Characterization of CHCs

In total, I identified 56 different CHC compounds of which eight were shared between *Scolopostethus pacificus* and *Liometopum occidentale* ; 15 compounds were unique to the bug, while 33 compounds were only detected in the ant (**Table 1**; **Fig 1**). The total amount of CHCs extracted from the bugs was higher (7.9±1.8 µg/mg; mean±SD) compared to the ants (4.2±3.7 µg/mg) (Mann-Whitney-U test: N=22, z= 2.34, p= 0.019, **Fig 1A**) and also the CHC composition was strikingly different (PERMANOVA: N_permutation_= 9999, F_1,20_= 1504, r^2^= 0.99, p< 0.001) showing no overlap in multivariate space (**Fig 1B**). The CHC profiles of *S. pacificus* mainly consisted of monomethyl alkanes with a C28 or C29 backbone chain length (**Fig 1C** - top). In strong contrast, CHC profiles *L. occidentale* were characterized by n-alkanes, a set of monomethyl alkanes and predominantly alkenes from the odd-numbered C25 to C31 series (**Fig 1C** - bottom).

**Table 1.**
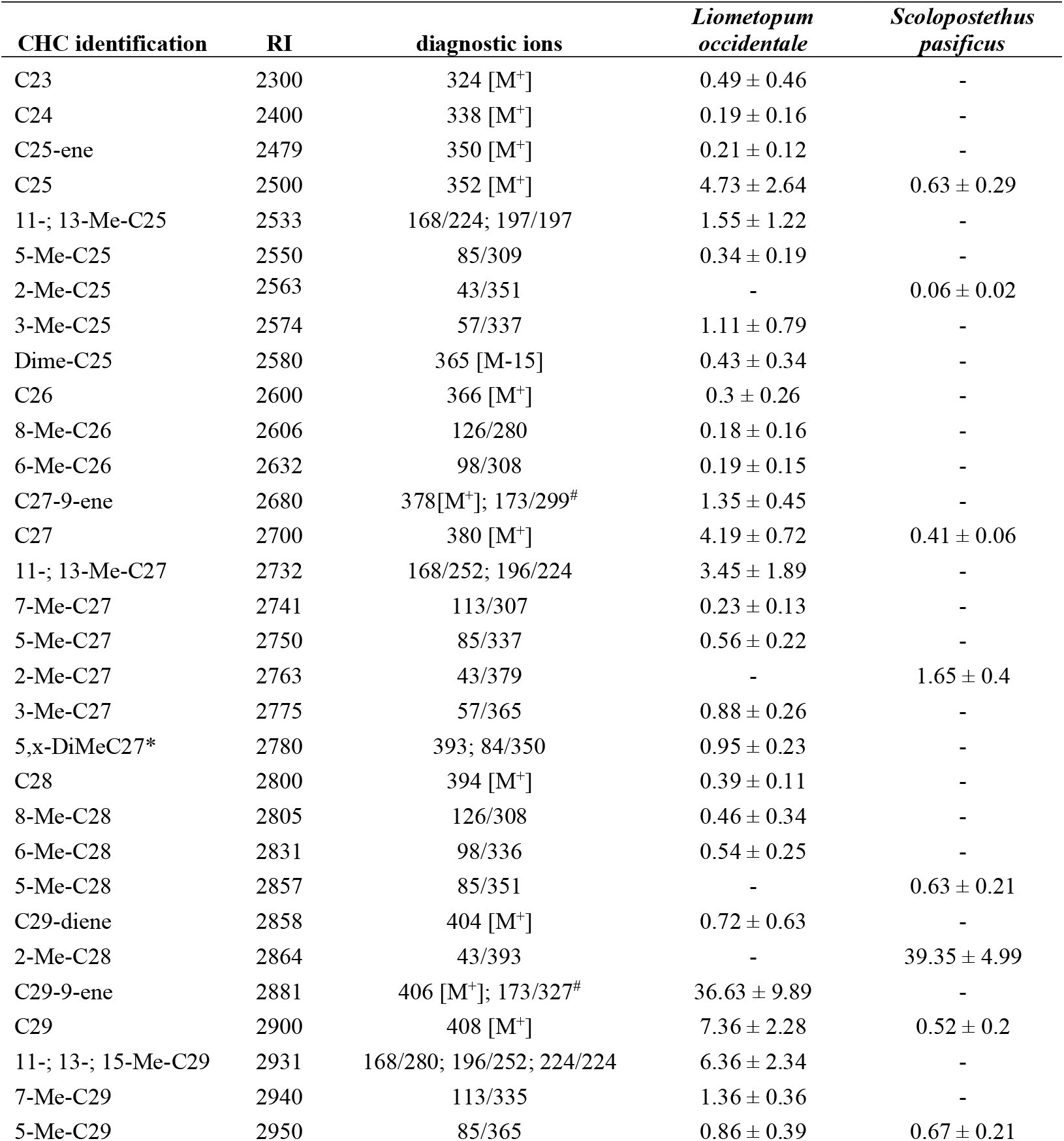
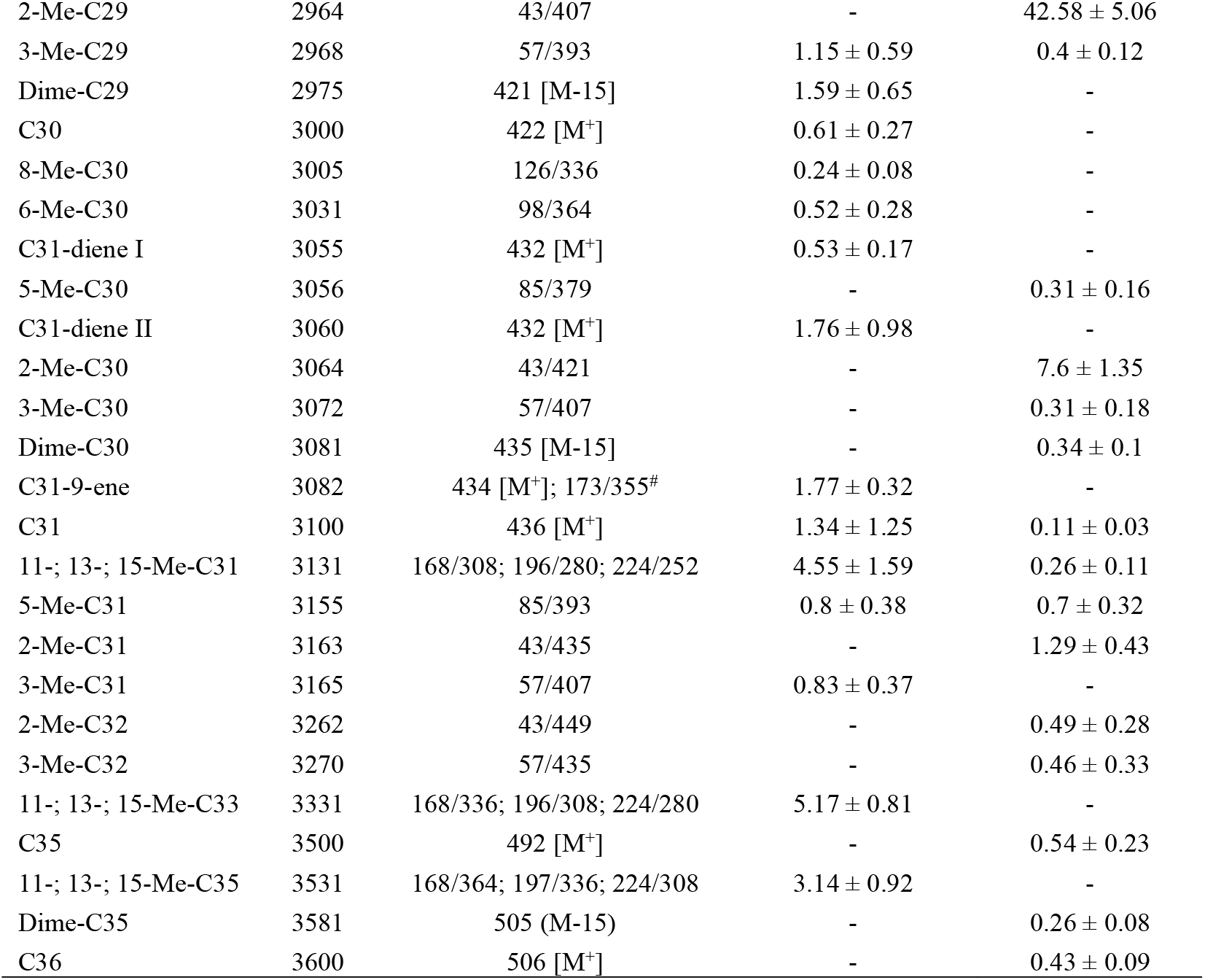
Cuticular hydrocarbons of *Liometopum occidentale and Scolopostethus pasificus*. Identified CHCs, their mean (±SD) abundances, the retention indices (RI) and diagnostic ions for identification. [M^+^] denotes molecular ions; diagnostic ions marked with # were obtained after DMDS derivatization of unsaturated hydrocarbons, *position of the second methyl branch unknown.

### Evidence for chemical defense

The five volatiles - (2*E*)-hex-2-enal, isoamyl acetate, 3-methyl-but-2-enyl acetate, (*E*)-4-oxohex-2-enal and (2*E*)-hex-2-enyl acetate (**Fig 2A; supplementary Table S2**) - were identified from the MTG of *S. pacificus*. The identity of four compounds, was confirmed with analytical standards and (*E*)-4-oxohex-2-enal was identified based on its *m/z* fragmentation patterns [112 (M^+^, 15), 97 (3), 84 (14), 83 (100), 57 (21), 55 (79), 53 (12); compared to Moreira and Millar (2005)] and retention index [RI= 960±2; RI_Lit_= 958; Yasuda et al. (2008)]. Fully-armed bugs had 1.3±0.3 µg of the volatiles mixture stored in their MTG, while bugs that were squeezed and washed to artificially disarm the insects had significantly less MTG content (0.3±0.2 µg; Mann-Whitney-U test: N=31, z= 4.72, p< 0.001; **Fig 2B**). Subjecting both armed and disarmed bugs to a 24h predation assay with ants, the overall survival of armed bugs (17/20 bugs alive) was much higher compared to disarmed bugs (1/20 bugs alive; GLM_binomial_: χ^2^= 15.16, N=40, p< 0.001; **Fig 2C**). Disarmed bugs that were kept in a similar setup for 24h without ants also showed a high survival (16/20 bugs alive, **supplementary Fig S2**). The 17 bugs surviving the bioassay had less secretion stored in their MTG compared to the armed insects (0.6±0.5 µg; Mann-Whitney-U test: N=33, z= 3.58, p< 0.001; **Fig 2B)**, indicating MTG usage against ants during the assay.

**Figure 2.**
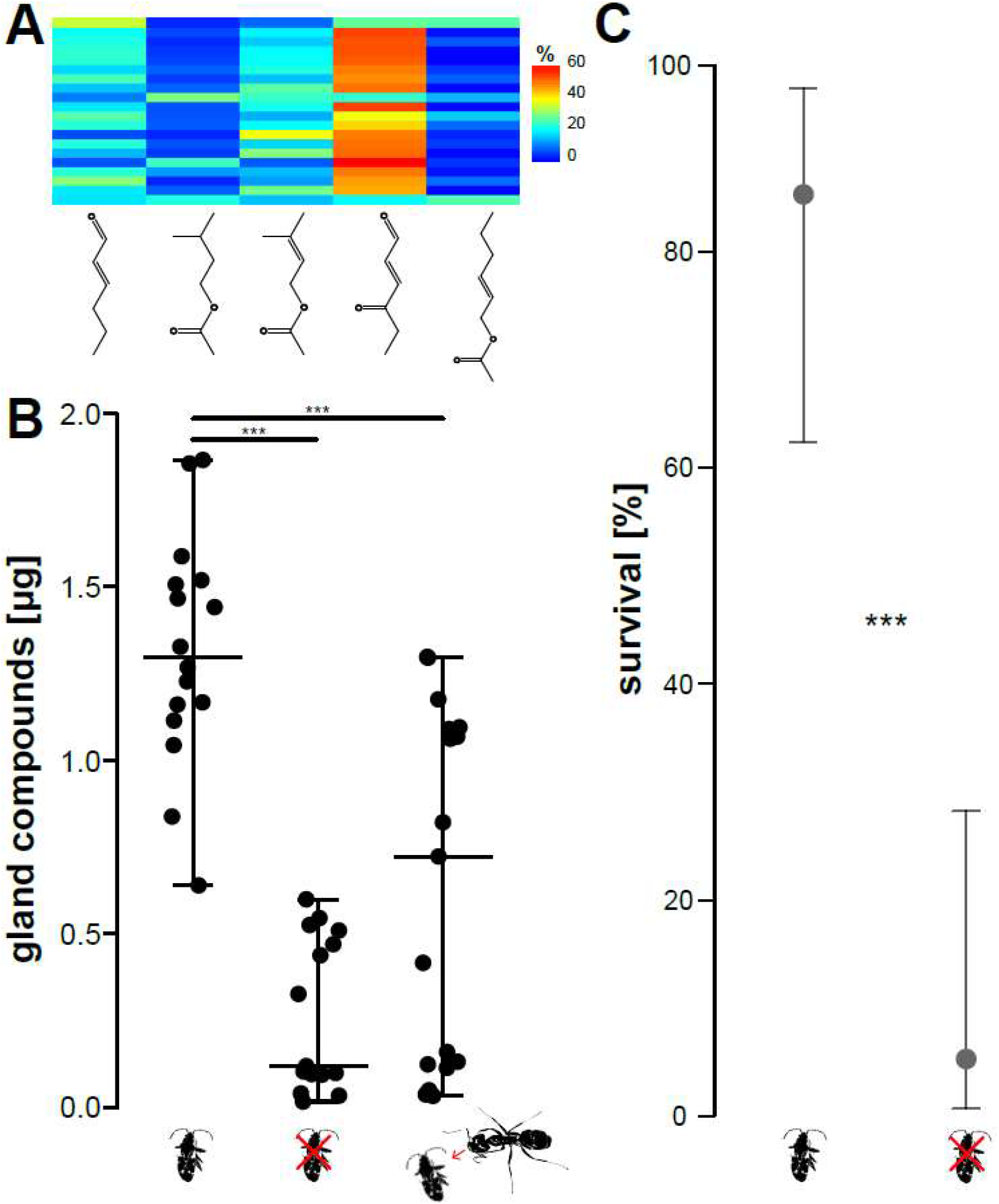
Chemical defense function of the bugs’ metathoracic gland. (A) The gland exudates of *Scolopostethus pacificus* are a blend of ketones, aldehydes and esters. The matrix plot shows the composition of twenty individuals. The gland compounds in the order of their retention indices are (2*E*)-hex-2-enal, isoamyl acetate, 3-methyl-but-2-enyl acetate, (*E*)-4-oxohex-2-enal and (2*E*)-hex-2-enyl acetate. The legend indicates the percentage composition for each compound per specimen. (B) Amount of MTG secretion [μg] of *S. pacificus*, in armed control bugs (normal bug pictogram), after artificially disarming via squeezing (red cross bug pictogram) and after bugs had been attacked by ants over a course of 24h (bug and ant pictogram). Horizontal bars represent the min, max and median; asterisks indicate significant differences (***= p<0.001) of Mann-Whitney-U tests. (C) Survival [%] of armed vs disarmed bugs after a 24h exposure to *Liometopum occidentale* ants. Dots depict the mean estimate and error bars are asymmetric standard errors of the GLM model (*** = p<0.001 of the GLM).

### Targeted molecular gut-content analysis

The ant-specific primers Loc1 (167 bp) and Loc2 (228 bp) showed amplification for ant DNA, but not for the bugs body (negative control) and guts (**Fig 3**). All replicates of the bug guts, bug bodies and ants showed amplification using the general ITS2 primers (**Fig 3**; **supplementary Fig S3A**). Guts dissected from the predaceous rove beetle *Platyusa sonomae* – which had been previously fed with *L. occidentale* – served as a positive control and showed PCR products for both Loc1 and Loc2 across all five positive control replicates (**supplementary Fig S3B**) and sequencing confirmed their identity (**supplementary text S1**).

**Figure 3.**
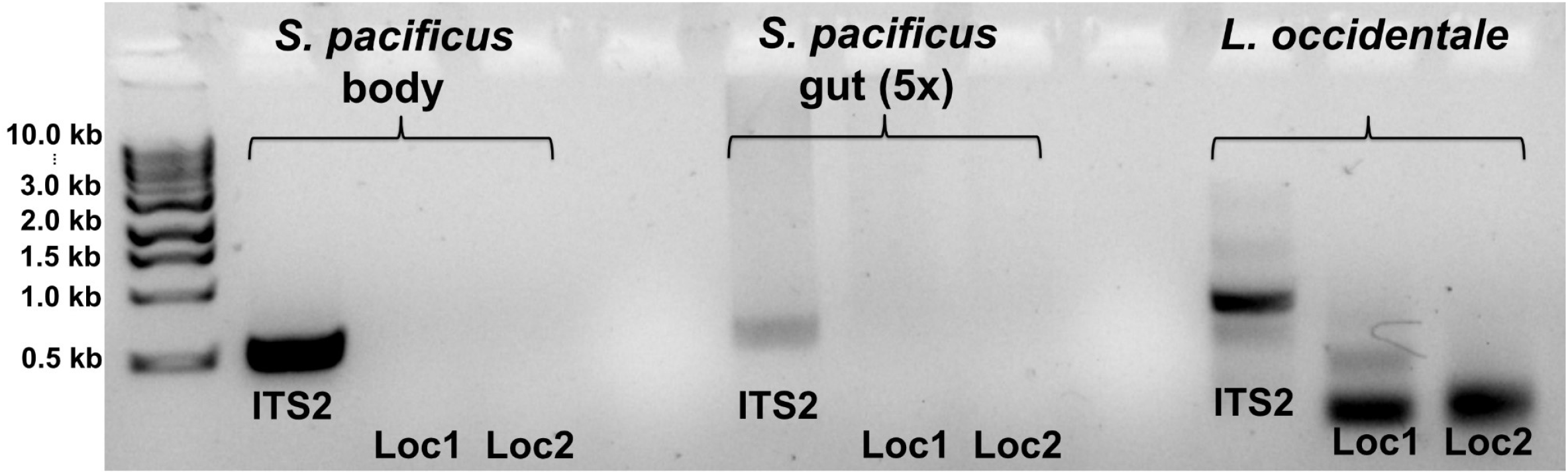
Molecular gut content analysis of *Scolopostethus pacificus*. Standard ITS2 (CAS5ps+CAS28s) and ant-specific Loc1 and Loc2 primers were used for PCR amplification using *S. pacificus* body-DNA, gut-DNA and *Liometopum occidentale* -DNA. While ITS2 amplified in all cases, no ITS2 fragment was amplified with ant-specific ITS (Loc 1/2) was amplified from dissected bugs’ guts.

## Discussion

Most studies on myrmecophiles have focused on well-integrated species and have deciphered their chemical integration strategies in great detail. While these animals represent remarkable examples of chemical trickery, as well as convergent and social evolution (Kronauer and Pierce 2011; Maruyama and Parker 2017; Parker 2016; Parker and Grimaldi 2014), there is also a diverse community of non-integrated ant associates (Parker 2016; Parmentier 2020; Parmentier et al. 2014). It has become clear that many such guests do not use any chemical resemblance strategies, are perceived as aliens and are attacked by the ants (Akino 2008; Gotwald Jr 1995; Kistner 1982; Parker 2016; Parmentier et al. 2017). These animals often escape their unwelcoming hosts by speed and agility or protect themselves with defensive chemicals or morphological structures (Akino 2008; Cushing 2012; Guillem et al. 2014; Parker 2016). In this study, I investigated the chemical strategies of such a non-integrated, facultative myrmecophile - the ant-resembling myrmecophilus bug *Scolopostethus pacificus*. I compared the CHC profiles of myrmecophile and host, analyzed the bugs’ MTG secretion and its function in ant-bug interactions as well as assessed aspects of *S. pacificus* feeding behavior. Chemical analysis revealed high amounts of CHCs in both species as well as distinct CHC profiles of the two species ruling out chemical mimicry and insignificance (**Fig 1**). Both the use of MTG secretions for defense (**Fig 2**) and the bugs feeding behavior (**Fig 3**) supported the idea that the parasite uses chemical weaponry to maintain an association with its host ants – presumably for protection.

The myrmecophilus bug *S. pacificus* had a CHC profile which clearly did not mimic the profiles of workers of its host ant species *Liometopum occidentale*. Additionally, the amount of hydrocarbons on the cuticle surface of the bugs and ants was in a comparable range. Therefore, *S. pacificus* clearly does not use chemical insignificance by reducing the CHC quantity – a strategy found in socially parasitic ant, e.g. *Ectatomma ruidum* or *Megalomyrmex symmetochus* (Akino 2008; Jeral et al. 1997; Neupert et al. 2018), but also in facultative, less integrated myrmecophiles (Parmentier et al. 2014; Parmentier et al. 2017). The CHC profiles of *S. pacificus* were dominated by monomethyl alkanes and n-alkanes while dimethyl-alkanes were only present in small amounts. The host *L. occidentale*, however, has CHCs mainly composed of C-29 compounds with very high proportions of alkenes (∼ 40%) and n-alkanes (∼ 18%), which is a common combination for ants (Kather and Martin 2015; Martin and Drijfhout 2009; Menzel et al. 2017). As methyl-branched compounds play an important role in recognition systems of many hymenopterans (Dani et al. 2001; Greene and Gordon 2007; Martin and Drijfhout 2009; Martin et al. 2011) - including ants - it comes as no surprise that *S. pacificus* is identified by the ants as an intruder and attacked, also ruling out “chemicals transparency” (i.e. use of compounds that are not recognized by the hosts; Martin et al. 2008).

The best-studied strategies used by parasites to overcome ant colony defense are CHC-based (Akino 2008; Bagnères and Lorenzi 2010; Lenoir et al. 2001), yet *S. pacificus* uses chemical weaponry to associate with its host ant. Bugs are well-known for releasing often-times malodourous volatiles from their MTGs when disturbed or threatened (Aldrich 1988; Krall et al. 1999). These volatile cocktails are typically composed of n-alkanes, alkenals and alkenyl alkanoates, but also monoterpenoid as well as aromatic compounds (Aldrich 1988). They usually function primarily as defensive chemicals to repel predators (Aldrich 1988; Krall et al. 1999). *Scolopostethus pacificus* is not different in this respect. Its MTG secretion is composed of well-known heteropteran volatiles that are expelled upon ant attack (**Fig 2B**). Using a manipulative approach of artificially disarming bugs, I was able to demonstrate that *S. pasificus* uses defensive chemicals against their host, enabling the bugs to survive in the presence of ants. This strategy might be comparable to the one of *Polistes* paper wasps, which secrete an ant-repellent chemical that hampers ants from invading the wasp nests and steal brood as well as stored food items (Post and Jeanne 1981). Unlike chemical weaponry in *Megalomyrmex* -fungus ant interactions (Neupert et al. 2018), however, *S. pacificus* was always attacked by *L. occidentale* (personal observation) resulting in high death rates of the myrmecophile in laboratory survival assays.

The targeted molecular gut-content analysis showed no sign of predation on ants in the bug. This is not surprising given that the vast majority of species in the dirt-colored seed bugs (Rhyparochromidae) are phytophagous, particularly feeding on fallen seeds (Schuh and Slater 1995). However, a single case of a zoophagus rhyparochromid is known from Ethiopia (Slater and Carayon 1963). Apparently, *S. pacificus* benefits from the association as it seeks close contact to the ant nests and as it invests into a costly defense (Thiel et al. 2018) against them. Furthermore, the bugs have evolved a visual appearance resembling their host ant. I consider it likely that the bug is a Batesian mimic of *L. occidentale*. While the adaptations to the associations with ants seem apparent, the benefits the bugs gain from this lifestyle are less clear. It may be that *S. pacificus* profits either directly from the enemy-free space, created by the aggressive and territorial velvety tree ants (Hoey-Chamberlain et al. 2013). These ants clear the nest perimeter and close-by areas from other predatory arthropods which could potentially prey upon the bug. More indirectly the bug might also exploit its cuticle patterns and color pallet that matches the one of *L. occidentale* very closely. This is very similar to myrmecomorph spiders – they resemble their host ant morphologically, adapt their behavior (i.e. erratic movement, akin to ants) and are thus considered Batesian mimics (Cushing 1997; Cushing 2012). These spiders are less likely to be chosen as prey by visually hunting predators that would otherwise eat them, conferring an adaptive advantage for the myrmecomorph mimics (Cushing 2012). That is why, the bug likely is a Batesian mimic as well, profiting from warning coloration shared with the velvety tree ant.

Overall, I showed that the ant-resembling bug *Scolopostethus pacificus* uses chemical defense based on its MTG to live in an association with *Liometopum occidentale* and is a potential Batesian mimic of the ant. In general, more research focusing on facultative myrmecophiles is needed to broadly understand the basic biology as well as the chemical and behavioral association mechanisms used by such ant guests.

## Acknowledgements

Thanks to Joe Parker and Mina Yousefelahiyeh for discussion and technical assistance, respectively. Steven Wilbert contributed pictures, Tom Naragon kindly provided *Platyusa sonomae* from his lab cultures and Betty Hong supplied analytical standards. Christiane Weirauch (UC Riverside) identified the bug species. Christoph von Beeren and Tom Naragon critically reviewed and commented on an earlier version of the manuscript. I am a Simons Fellow of the Life Sciences Research Foundation (LSRF).

## Ethics statement

There are no legal restrictions on working with the herein mentioned species. Field collection permissions were issues by California Department of Fish and Wildlife and the Angeles National Forest (US Forest Service; USDA).

---

**Supplementary text S1** ITS2 nucleotide sequences of the ants, bugs and *Platyusa* beetles as assessed by Sanger sequencing as well as sequenced PCR products of Loc1 and Loc2 from the positive control.

`>ITS_2_Liometopum_occidentale`

TCAGCGGGTAATCTCGCCTGCTCTGAGGTCGTCGAATACAGAATAAAAAAATTCCCCTCTTTTTCTCTCTTCTACCCGCTGCTTCTCTCTCTCTCTCTACTCGTCGCCGCAGTCGTCGTGCGTGACGACGCGCGTCGAAAGAGTGAGCGCCGAGAGAGCGAGGTAGAGACGAGAAAGGAGGAGGAGGTTCTCGCACGTGTGTATCGGGGTGTTGATAATCCAGACGCGATCGCACGACACACGACATGCCGGTCCTTAGAGCTGGGCCGCAATAGGAAGACTCCTCTTCTCTCCTCTGACACGTCGCGCTCTCTGACTGAGAGAGACGCAACGGCGAGGAGAGAGAGAGAGAGCTCCTTTGAGGCCTGCGAAATTCCGTGGACTCGGACACGCGGTCGGCTCGCGCGGAAAAACCCACCACCGATTCGTAACCGTGGCACAACGTATCGTATGTGTATTTACTGGAACACATCGCGACACGCCACGATACGCGCGCGCTCGTGCCCCTCGGAGGTGTGCCTCGGAAGGGAAGACTGAATGAGAGAATGCCGGCTTTTCTCTCTCGCCACCGTACGAGATGTCGGGTGCGTAACGGGGACGCACCGTGGTCGACGACATATCTCCATCGGTCGCGAAAACGCCGATTCTCTCTCTCGTCTCGCACACATTCCGCGCCACCTCCTTGGCGGCCCCAAGGGCCGCGGCGGACACGCGCGCGAACACGCGCTCTCTCCGTGTCATTTCAGACGACGCCCAGGGGGGCGAGACCTCTCACTCCTCGTCGGTGTGTGTGGTAACGAAGACCGCACTCCGCGTGAGGAGCGTCTCGCGCGCGCAACCTCGGCGAACGTCCAACGTCGCTCGTACGCGCGCGTCTCGACGACACGCGCGCAAGCAGTTTCGGGGTACTGAAACGACCCTCAGCCAGGCGTGGTCCGGGAATGT

`>ITS_2_Scolopostethus_pacificus`

ATTTTGAACGCACATTGCGGCCCTGGGCTAGAACCAAGGGCCACCCCTGTCTGAGGGTCGTTTGTACTAAAAGGAGACTCTCGAGTCTCGAAAAACGATGTCTCTTTAACTTCCTGTCGCCCTTTGCGGCTTGACTGGAGGGAGACGAGAGGGGAGTTCCGCTCGCCAGTCTCCGTGAGGTGACAAAGGCGACCGCCTCCCTACAAAAGCCAATGCTCTGCCGTAAGCCGAGGCGCTCGACGGACTCTGGTCTTCCCCAACCGCGAGGTAGAGGAAGGCTACAACCGTACACCTGGTTCTTCGAAAGGGGTAAATTTTTTTCTCCTCTCTCCGGCCGAGCACGACCCGTTGTTGGTTGATACCCGAGCGGGCTCGACAGGTTCGGATTCGAGAGGAGGAAAACTTTCCCCTTGCATCCTCGAAGGTACCAACGTTACCGGTTTTGAAAGGGTTCGTCAAGAGCAAGCCAAGCTTAACAAAACGGCGCTATCATTATTTTCGACCTCAGATCAGGTGGGGCTCCCCGCCGAATTTAAGCATATCAATAAGCGGAGGAAAAGAAA

`>ITS_2_Platyusa_sonomae`

ATATGCTTAAATTCGGCGGGTAGTCTCACCTGCTCTGAGGTCGCGTATTTGAGAGTTATACTAAAAGTATGCATCTCGTGGTTTCATGTCAAATTACGTAGTTCGGTTATGCTTGTCGTGATTGAAATGCGAAAACACCTTAACTTATGCGGTATTGCTCAGCAAGCGCGTCGTGCCGCCGCCGTACTGGGACAAGCTACCCGTCGTGACTCGACATTGTCGTCATGGACGCGACGGACTTGGCCCAACGTATATGCGGACGATCGCTGCGTTAGCAAGCGCAAGGTATAAACCGAAAACGCTACACAATACACTGTGATAACCAAGCCTGACATTGAAAATCAATCGCGTAATTCCCTATCGACTAACGCCGAATCGGGCACGATCGAGTATTTAAAGAGACGAACAACGTGCGAAACGTTGCGCGAGCTCCCAATTCGTCCCGCAATTAACCACGTGCAATACGAGTTAATCCGGACAGTCGTATAAATTGAAACGACCCTCAGCCAGGAGTGGTCCAGGAACAGTATCCAAGGACCGCAATGTGCGTTC

`>Loc1_Platyusa_PCR_product`

NNNCNNGGNNNNANNNNGGCCACCCCTGAGGTCAGACGAGAAAGGAGGAGGAGGTTCTCGCACGTGTGTATCGGGGT GTTGATAATCCAGACGCGATCGCACGTCACACGACATGCCGGTCCTTAGAGCTGGGCCGCAATAGGAAGACTCCTCT TCTGTCCTCTGACACGTCGCGCTCTCTGACTGAGAGAGACGCAACGGGCCACCCCTGAGGTCNNNCNNGGNNNNANN NN

`>Loc2_Platyusa_PCR_product`

NNAAATAATGGGTAGTCTNNNNNNGAATAATGATCTGACTGAGAGAGACGCAACGGCGAGGAGAGAGAGAGAGAGCTCCTATGAGGCCTGCGAAATTCCGTGGACTCGGACACGCGGTCGGCTCGCGCGGAAAAACCCACCACCGATTCGTAACCGTGGCACAACGTATCTTATGTGTATTTACTGGAACACATCGCGACACGCCACGATACGCGCGCGCTCGTGCCCCTCGGAGGTGTGCCTCGGAAGGGAAGAGGGTGCAATANNNANNNATGTTCAANN

## Supplementary Tables

**Table S1.**
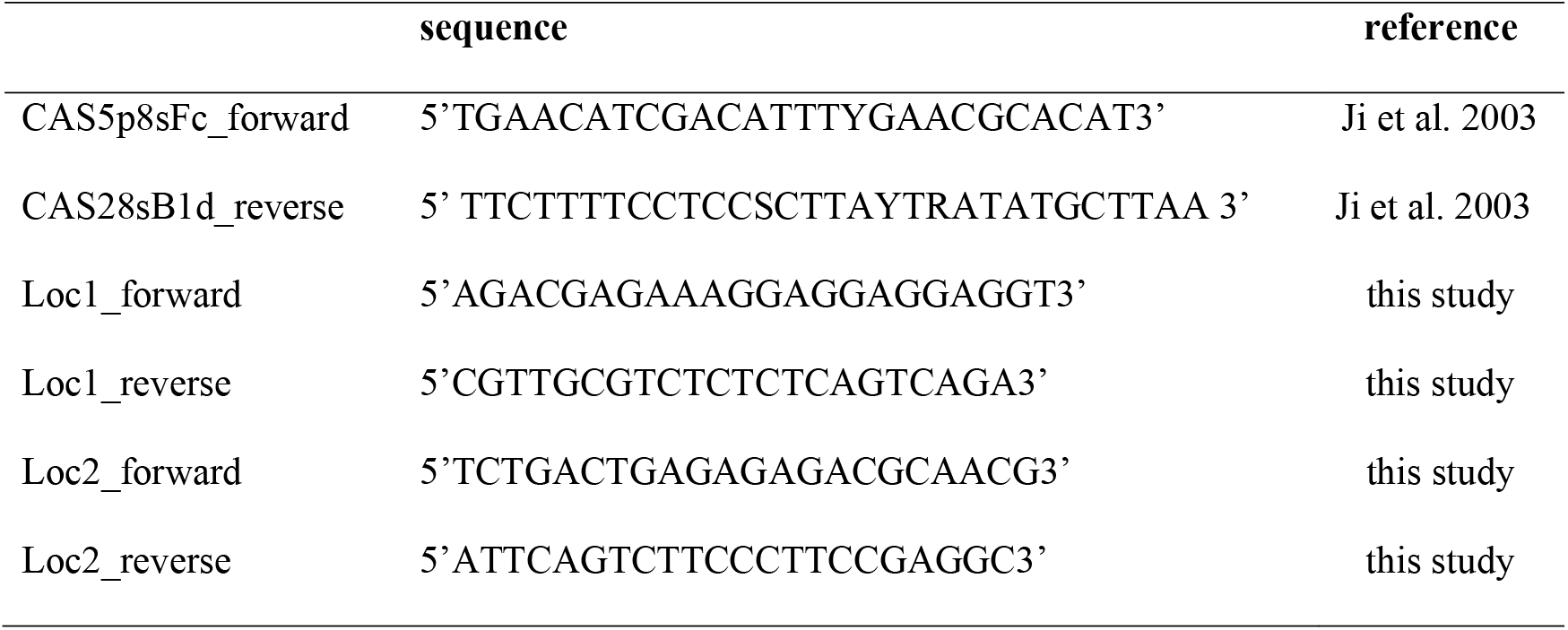
Primers sequences used in this study.

**Table S2.**
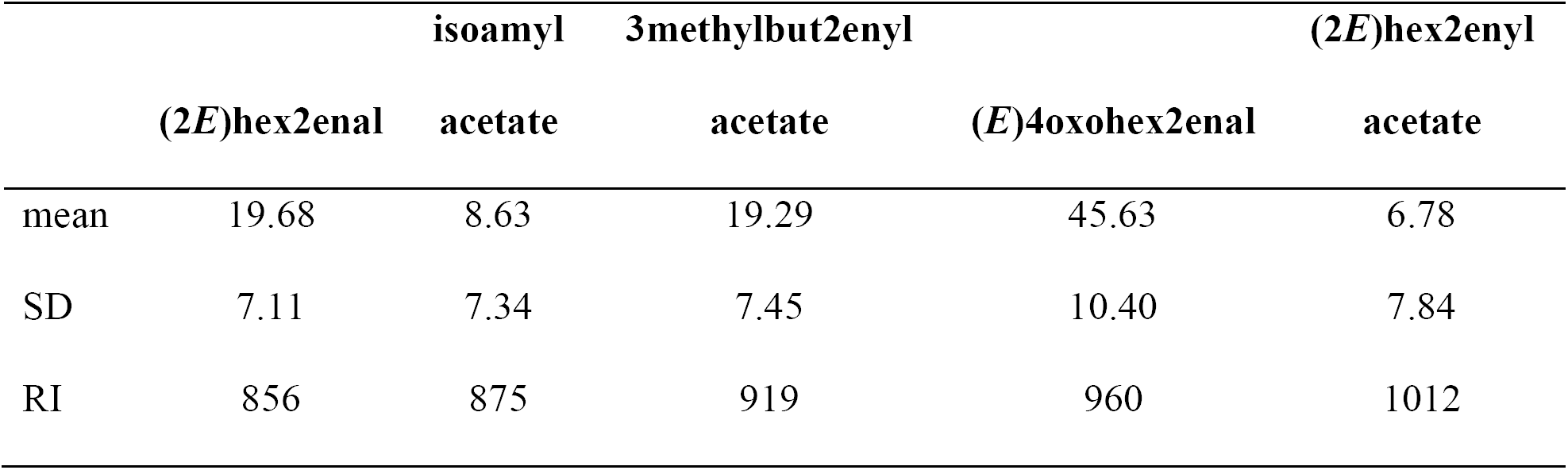
Proportions (mean ± SD) and retention index (RI) of volatile MTG compounds from *Scolopostethus pacificus* identified via SPMEGC/MS and whole body extract GC/MS

## Supplementary Figures

**Figure S1.**
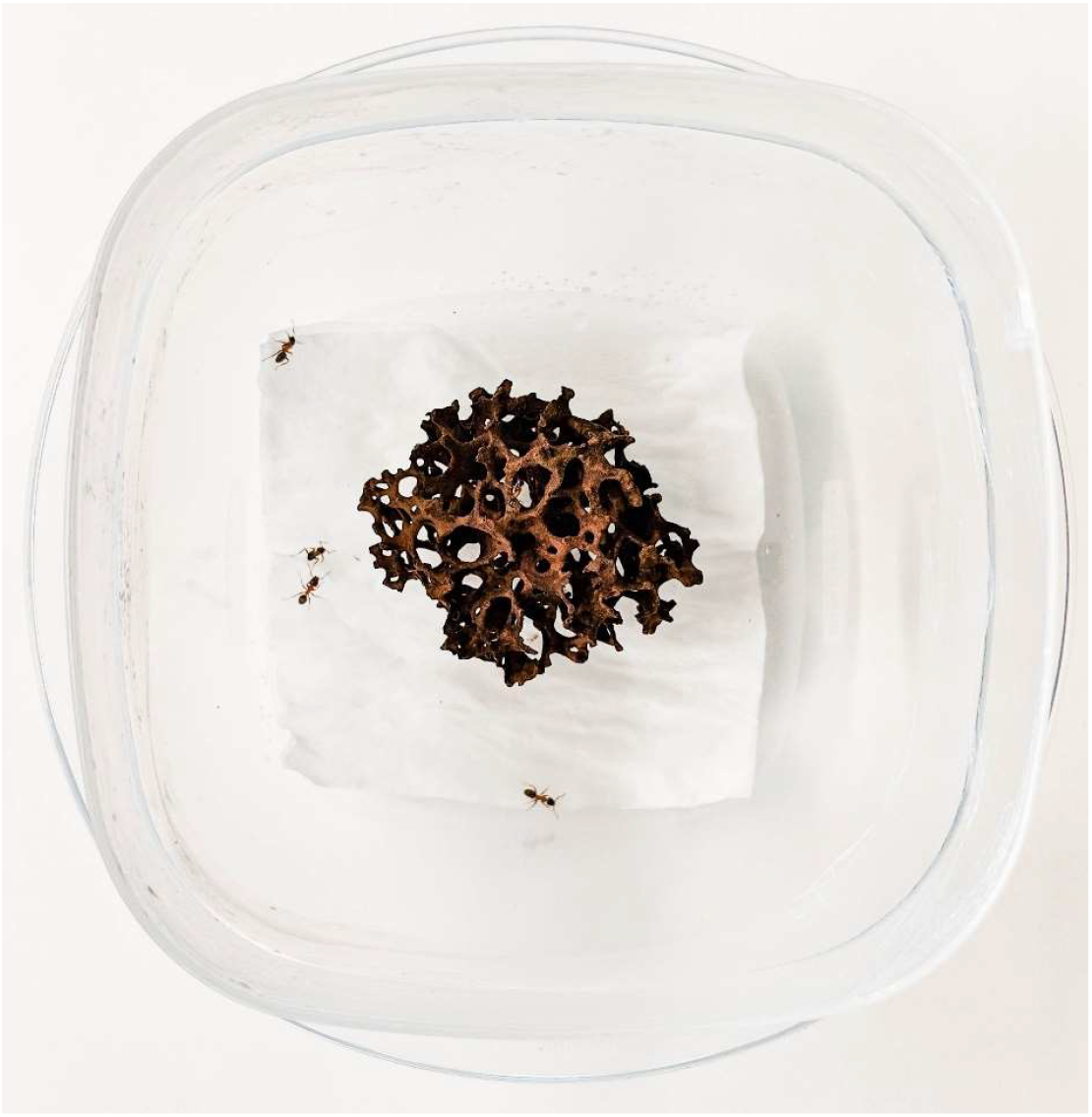
Setup for the survival bioassay trials: five *Liometopum occidentale* were put into small plastic boxes (105×105×80 mm) with a piece of nest material and the bottom was covered with wet tissue paper. One *Scolopostethus pacificus* bug (from different treatments) was added per box and its survival assessed after 24h.

**Figure S2.**
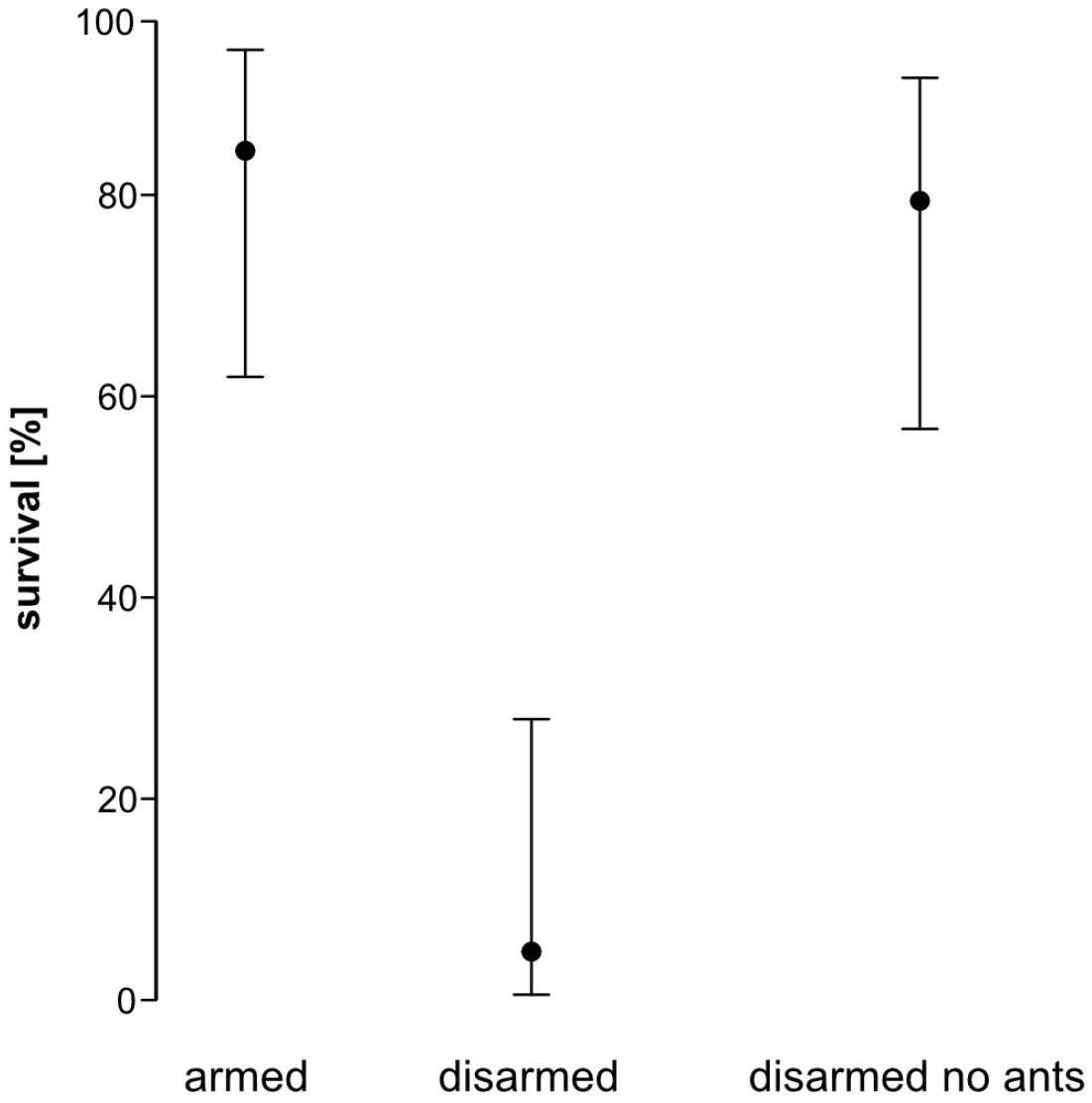
Survival [%] after 24h of armed bugs vs disarmed bugs exposed to *Liometopum occidentale* ants vs disarmed bugs with no ants. Dots depict the mean estimate and error bars are asymmetric standard errors of the GLM model.

**Figure S3.**
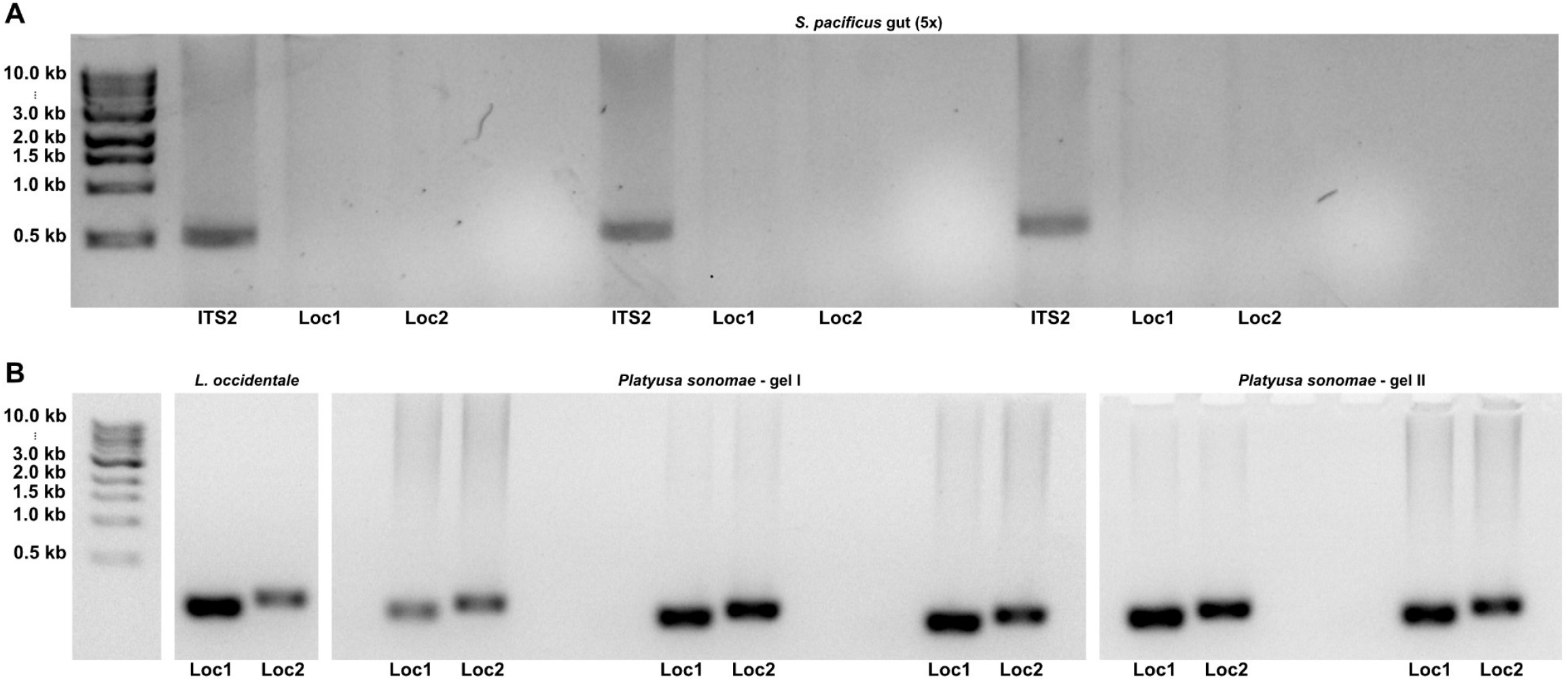
(A) Molecular gut content analysis of *Scolopostethus pacificus*. Standard ITS2 primers (CAS5ps+CAS28s) amplified the 563bp long bug ITS2, but no ITS amplification was detected with ant-specific Loc1 and Loc2 primers. (B) As a positive control the ant-specific primers Loc1 (167 bp) and Loc2 (228 bp) were amplified with ant DNA. To demonstrate that Loc1 and Loc2 can amplify ant DNA from a predator which recently consumed ants, I dissected guts of *Platyusa sonomea* rove beetles, that had preyed on ants. For all give rove beetle samples, Loc1 and Loc2 amplified and their identities were confirmed with Sanger sequencing. The ladder shown on the left (A+B) is the 1 kb DNA Ladder from New England Biolabs.

